# Endothelial Cell Secretome Alterations Induced by Inflammatory Stress

**DOI:** 10.64898/2026.01.15.695726

**Authors:** Vi T. Tang, Srishti Baid, Alejandra Castellanos, Venkatesha Basrur, Colin A. Kretz, Alexey I. Nesvizhskii, Xiang Gao, Qunfeng Dong, Mitchell J Weiss, Prabhodh S. Abbineni

## Abstract

Endothelial cells detect pathogens through pattern recognition receptors, such as Toll-like receptor 4 (TLR4), which triggers the synthesis and secretion of molecules that initiate the innate immune response. Proteins bearing signal peptides are secreted through the classical endoplasmic reticulum (ER)–Golgi–dependent route, whereas select signal-peptide-lacking cytoplasmic proteins are secreted via less well-characterized ER-Golgi-independent mechanisms, collectively termed unconventional cytoplasmic protein secretion (UCPS). To systematically characterize the secretome of human umbilical vein-derived endothelial cells (HUVECs) and delineate the contribution of UCPS, we performed deep quantitative proteomics on HUVEC cell lysates and conditioned medium before and after TLR4 stimulation with lipopolysaccharide (LPS). Of 5205 proteins detected in either fraction, 381 were enriched in the conditioned medium and therefore classified as secreted. Of these, 333 proteins (87.4%) were secreted via the conventional pathway, and 48 (12.6%) were secreted via UCPS, 43 of which were not previously associated with this process. Predicted functions of UCPS-secreted proteins include redox regulation, proteostasis, cytoskeletal remodeling, and innate immune signaling. We confirmed that α-globin (HBA1), which functions as a redox sensor and regulator of nitric oxide in endothelial cells, is secreted constitutively by UCPS and at higher levels following inflammatory activation. Notably, UCPS cargo identity showed poor concordance with current computational predictors, underscoring the need for empirical datasets. Overall, our findings suggest that the HUVEC secretome includes both conventionally and unconventionally secreted proteins that regulate coagulation, angiogenesis, and immune function. Our findings establish a high-quality secretome dataset for HUVECs, providing a novel resource for future efforts to define the molecular determinants governing UCPS cargo selection and trafficking related to endothelial cell function.

## Introduction

Endothelial cells regulate vascular function and play an active role in innate immunity by detecting inflammatory signals and coordinating immune responses^1-3^. Endothelial cells sense and respond to inflammatory stimuli via pathogen and damage recognition receptors, including the toll like receptor (TLR) ^4-8^ and the nod-like receptor (NLR) ^9-11^ families. Upon activation, endothelial cells synthesize and secrete numerous proteins that initiate and amplify the inflammatory immune response. These include pro-inflammatory cytokines and chemokines that recruit and activate innate and adaptive immune cells, as well as adhesion molecules that facilitate immune cell retention at inflamed sites^12, 13^.

Protein secretion in endothelial cells occurs through both conventional and unconventional secretory pathways ^14-16^. Signal-peptide bearing proteins are co-translationally inserted into the endoplasmic reticulum (ER), trafficked to the Golgi^17, 18^, and then either constitutively secreted via vesicle fusion with the plasma membrane, or in the case of von Willebrand factor (VWF) and a few other molecules, stored in post-Golgi secretory organelles known as Weibel Palade bodies^19,20^. Additionally, certain cytoplasmic proteins lacking signal peptides are secreted via ER-Golgi-independent routes, termed unconventional cytoplasmic protein secretion (UCPS) ^16, 21-23^. While the conventional, signal peptide-dependent, secretory pathways of endothelial cells are well-characterized, cytoplasmic cargoes secreted via the UCPS are poorly defined^24, 25^. Furthermore, the extent to which inflammatory signals modulate the endothelial cell secretome, particularly in the context of UCPS, remains poorly characterized.

We analyzed inflammation-induced alterations in the secretome of primary human umbilical vein-derived endothelial cells (HUVECs) using a tandem-mass-tag (TMT)-based quantitative proteomics approach that effectively distinguishes secreted from intracellular proteins by assessing their relative abundance in conditioned medium and cell lysates^24, 26^. As demonstrated previously, the media-to-lysate abundance ratios provides a practical approach to distinguish secreted proteins from background contamination^25-29^. This enabled the identification of both conventionally and unconventionally secreted proteins and allowed us to determine how lipopolysaccharide (LPS), a bacterial endotoxin, reshapes the endothelial secretome. While most secreted proteins from HUVECs utilize the ER-Golgi pathway under resting and inflammatory conditions, we found evidence for UCPS and identified novel cargoes secreted via this pathway. We validated unconventional secretion of α-globin, which is secreted by resting cells and at higher levels following exposure to inflammatory stimuli.

## Experimental Procedures

### Cell Culture

HUVEC (Lonza Bioscience) were plated on tissue culture dishes coated with 2% collagen, and cultured in EGMTM-2 Endothelial Cell Growth Medium-2 (Lonza Bioscience) which contains the growth factors VEGF, hFGF, IGF, EGF, and 2% fetal bovine serum. HUVEC were expanded until they reached ∼80% confluence in 10-cm dishes and used for proteome and secretome analysis described below.

### Conditioned Medium and Cell Lysate Collection

For secretome analysis, cells were washed three times with pre-warmed PBS and incubated in serum-free, phenol red-free DMEM for 4 hours (10 ml per dish). For brefeldin A (BFA) treatment, cells were first pre-incubated with serum-free DMEM containing 1 µg/ml BFA (or vehicle control) for 30 minutes. The medium was then discarded and replaced with fresh medium containing 1 µg/ml BFA, followed by incubation at 37°C for an additional 4 hours. To examine the influence of inflammatory stress on the endothelial secretome, HUVECs were treated with 1 µg/mL lipopolysaccharide (LPS). In one condition, cells were incubated with LPS for 4 hours in serum-free DMEM (Thermo Fisher Scientific, Waltham, MA, 31053-036). Alternatively, HUVECs were pretreated with 1 µg/mL LPS for 20 hours in complete medium, followed by a 4-hour incubation in serum-free DMEM. In select experiments, 10 µM nigericin was added during the final 45 minutes of the 4-hour serum-free incubation, either alone or in combination with prior 20-hour LPS priming.

Conditioned medium was collected by transferring the medium to 15-ml conical tubes and processing all subsequent steps at 4°C. The medium was centrifuged at 500 × g for 5 minutes to remove cellular debris, followed by a second centrifugation at 5000 × g for 20 minutes. The clarified medium was concentrated using Amicon ultrafilters with a 3 kDa molecular weight cutoff (Thermo Fisher Scientific, Waltham, MA, 31053-036). Approximately 10 ml of medium was concentrated to 200 µl by centrifugation at 4000 × g for 40 minutes. Protein concentration was determined using DC protein assay (Bio-Rad, Hercules, CA, 500-011), and samples were aliquoted and stored at –80°C. For lysate preparation, lysis buffer was prepared by supplementing 10 ml of RIPA buffer with a protease inhibitor cocktail (cOmplete Mini Protease Inhibitor Cocktail, Roche, Basel, Switzerland, 11836153001). After removal of the medium, cells were washed once with PBS. A total of 500 µl of lysis buffer was added per dish, and the cell suspension was collected into microcentrifuge tubes. Samples were incubated on ice for 30 minutes, briefly sonicated, and centrifuged at 20,000 x g for 45 minutes at 4°C to pellet insoluble material. The resulting lysate was transferred to new microcentrifuge tubes, protein concentration was measured, and samples were aliquoted and stored at –80°C.

### Mass Spectrometry

Mass spectrometry-based protein identification and quantification were performed as previously described. Briefly, 50–75 μg of total protein from each cell lysate and conditioned medium sample was subjected to proteolytic digestion using a trypsin/Lys-C mix (1:25 protease:protein; Promega) and labeled with either TMT 10-plex or TMT 16-plex reagents (Thermo Fisher Scientific, 90110), following the manufacturer’s protocol. Labeled peptides were analyzed by liquid chromatography–tandem mass spectrometry (LC-MS/MS), as previously described^24, 26^. Raw data files were converted to mzML format and processed using the FragPipe computational pipeline (https://fragpipe.nesvilab.org/) with the default TMT10-MS3 workflow. An adjusted *p*-value (*q*-value) of 0.05 or less was considered statistically significant.

### Immunoblotting

For immunoblotting analysis, 15 ug of lysate and 20 ug conditioned medium protein were resolved using SDS-PAGE (Invitrogen XP04205BOX), followed by transfer onto nitrocellulose membrane. Following transfer, nitrocellulose membranes were incubated in in 1X Tris Buffered Saline (TBS, Biorad 1706435) containing 0.1% Tween (Sigma P9416-100ML) and 5% milk for 2 hours, followed by 3 washes in TBS containing 0.1% Tween. The primary antibodies rabbit polyclonal Hba1 (1:300 dilution), interleukin 8 (1:1000 dilution, R & D system MAB208-100), and beta-actin (1:1000 dilution Santa Cruz Biotechnology, sc-47778) were added overnight.

Nitrocellulose membranes were then rinsed and incubated with either goat anti- rabbit HRP (Biorad,170-6515) or mouse anti-rabbit HRP (Biorad,170-6516) at a 1:5000 dilution. The chemiluminescent signals were assessed following exposure to SuperSignal West Femto Maximum Sensitivity Substrate (Thermo Fisher, 34095).

### Experimental Design and Statistical Rationale

For all proteomic experiments, HUVECs derived from pooled donors were expanded under identical culture conditions and plated in matched 10-cm dishes. Each experimental comparison was performed using three independent biological replicates, where each replicate consisted of a distinct dish processed from separate expansions of the pooled-donor HUVEC population. For every replicate, matched cell lysate and conditioned medium samples were collected from the same dish, enabling paired calculation of medium-to-lysate (M/L) ratios.

For proteins detected in both lysate and medium fractions, a medium-to-lysate (M/L) ratio was calculated based on absolute reporter ion intensities. Signal peptide and transmembrane domain annotations were retrieved from the UniProt database. The limma statistical package was used for comparison of protein abundance or M/L ratios between various conditions described in the results section, and p-values were adjusted for multiple hypothesis testing using the Benjamini and Hochberg method^30^. M/L ratios were always calculated from strictly paired lysate and medium samples obtained from the same dish within the same plex. An adjusted *p*-value (*q*-value) of 0.05 or less was considered statistically significant.

The HUVEC mass spectrometry data was used to assess the accuracy of SecretomeP and Outcyte in predicting unconventional protein secretion. Two protein groups were taken from the secretome data: (1) a "media only" group of 38 proteins signal-peptide lacking, brefeldin-a resistant protein detected in the conditioned medium, and (2) a group of 50 proteins predominantly found in the cell lysate, with a M/L ratio<1 which was used as a negative control. UniProt was used to obtain amino acid sequences of proteins in both groups. Fasta files were uploaded to SecretomeP and Outcyte and each tool generated UCPS prediction scores, which were plotted using Prism 10. Statistical analysis was performed using an unpaired t-test. Statistical significance is indicated as: * (p< 0.05), ** (p< 0.01), *** (p<0.001).

## Results

### The Endothelial Secretome Is Composed of Proteins Secreted via Conventional and Unconventional Secretory Pathways

We previously validated a mass-spectrometry based secretome approach that measures protein abundance in both conditioned medium and cellular lysate fractions to calculate a medium-to-lysate protein abundance ratio (M/L ratio), and showed that high M/L ratios indicate secreted proteins ^24, 26^ (**Fig. 1A**). This approach identified secreted proteins in HUVECs, as evidenced by an increase in the percentage of proteins with signal peptides correlating with higher M/L ratios (**Fig. 1B**). All proteins with M/L ratios > 10 contained signal peptides, indicating that the most robustly secreted proteins in resting HUVECs utilize the conventional ER-Golgi secretory pathway (**Fig. 1B**). GO term enrichment of this group identified functions such as regulation of blood coagulation, positive regulation of leukocyte chemotaxis, and angiogenesis, all known to be regulated by endothelial secretion (**Supplementary File 1**). In the intermediate M/L ratio group (1-10), 24% (25 proteins) lacked signal peptides, suggesting they represent UCPS cargoes (**Fig. 1B**). Additionally, 252 proteins were detected solely in the medium fraction, precluding M/L ratio calculation. Of these, 15% (38 proteins) lacked signal peptides and transmembrane domains, nominating them as potential UCPS cargoes (**Fig. 1B**).

**Figure 1.**
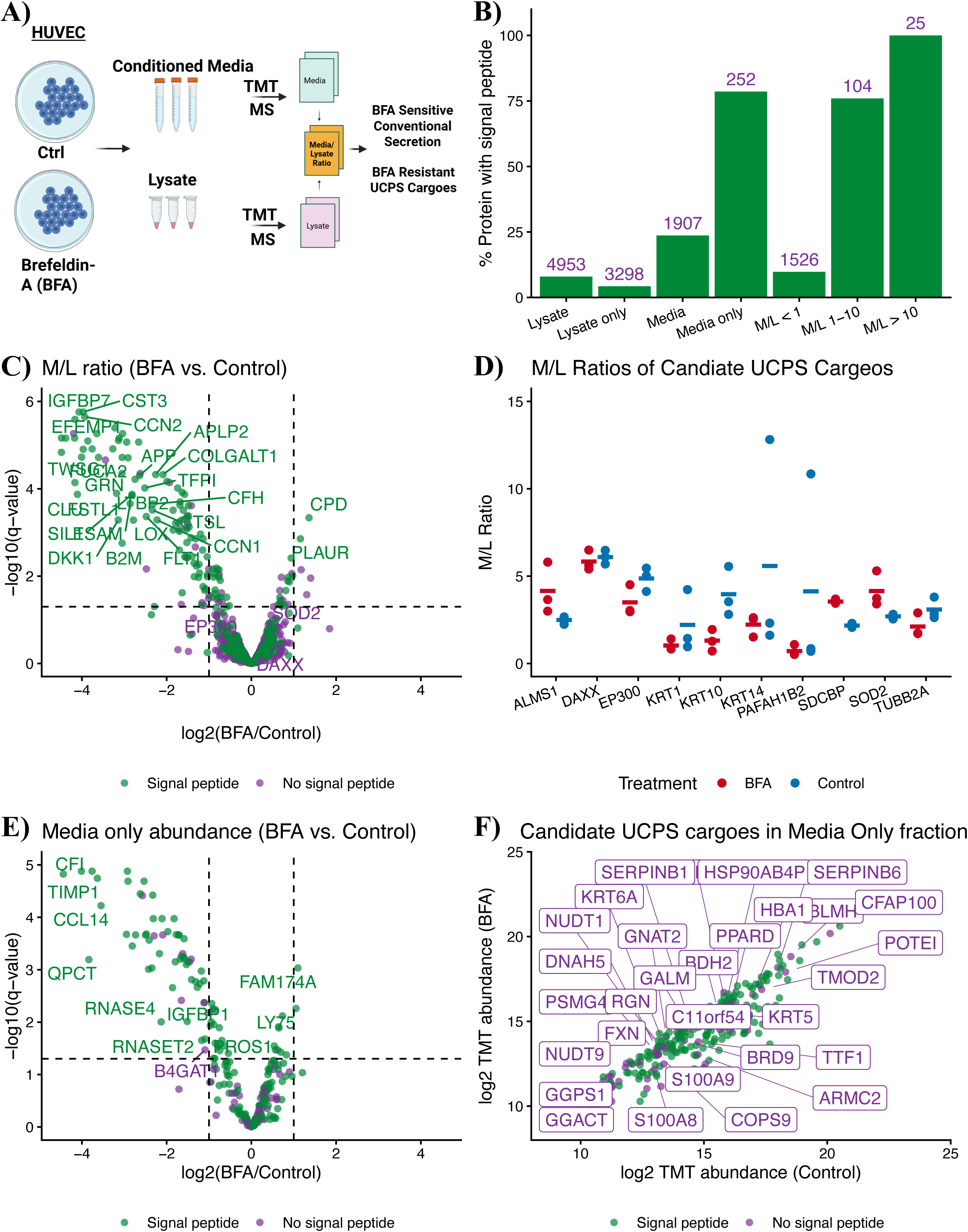
Identification of BFA-sensitive conventionally secreted proteins and BFA-resistant UCPS cargoes secreted by HUVEC. **A)** Schematic illustrating approach for calculation of media/lysate protein abundance ratios (M/L Ratio). **B)** Percentage of proteins harboring signal-peptides in media and lysate fractions, and M/L Ratio groups (n = 3). The number of proteins per group is indicated above each bar. **C)** Volcano plot comparing M/L Ratios of brefeldin-A treated versus control HUVEC (n = 3 per group). **D)** Candidate brefeldin-A resistant UCPS cargoes (n = 3 per group). **E)** Volcano plot comparing abundance of proteins identified in the Media Only fraction in brefeldin-A treated cells versus control HUVEC (n = 3 per group). **F)** Candidate brefeldin-A resistant UCPS cargoes identified in the Media Only fraction (n = 3 per group).

To confirm that UCPS cargoes use ER-Golgi independent routes, we treated HUVECs with one μg/ml brefeldin-A (BFA), an inhibitor of ER-Golgi transport that selectively blocks conventional secretion. Upon treatment with BFA, there was a significant accumulation of signal-peptide containing proteins in the cell lysate (**Fig. S1A**), depletion from the conditioned medium (**Fig. S1B**), and a consequent reduction in the M/L of ratios of 109 proteins (**Fig. 1C**; green and purple datapoints indicate proteins containing and lacking signal peptides, respectively). Of these, 92.6% had signal peptides or transmembrane domains, indicating secretion via the ER-Golgi pathway. The remaining 8 proteins (ANKFY1, ATP5IF1, CTTNBP2NL, KRT10, MAP2K1, PPA2, PPIC, RPA2) did not contain either signal peptides or transmembrane domains, but were sensitive to BFA treatment, indicating either cryptic, unannotated signal peptides, or indirect effects of BFA treatment. In addition, we identified candidate UCPS cargoes as those that lack signal peptides exhibit high M/L ratios (M/L 1-10) that are resistant to treatment with BFA (**Fig.1D**; Blue and red datapoints represent control and BFA treated conditions, respectively). As expected, these proteins display unchanged log2FC after BFA treatment relative to control and therefore cluster around the log2FC ∼ 0 region in the volcano plot shown in **Fig. 1C**. This included the known UCPS cargoes superoxide dismutase 2 (SOD2) and syntenin-1 (SDCBP), and several novel candidates (**Fig. 1D**).

BFA treatment reduced the abundance of signal-peptide or transmembrane domain-containing proteins detected solely in the conditioned medium fraction (**Fig. 1E**). 38 proteins (15%) detected in this fraction lacked both signal peptides and transmembrane domains, and were resistant to treatment with BFA, making them candidate UCPS cargoes (**Fig. 1F**). Of these, S100A8 and S100A9 are known UCPS cargoes that induce neutrophil chemotaxis and adhesion^31^. This fraction also contained several novel UCPS cargoes. For example, serpinb1 and serpinb6, members of the serine protease inhibitor B family, have known intracellular roles in regulating inflammatory protease activity^32-34^. One study demonstrated that pancreatic beta cells secrete serpinB1 ^35^, but it’s extracellular function and protease targets are not well defined. Beyond these, the UCPS cohort included redox enzymes (e.g., BDH2^36^, HBA1 ^37^), proteostasis factors (e.g., BLMH^38^, COPS9 ^39^), and cytoskeletal components (e.g., TMOD2 ^40^), implicating endothelial UCPS in stress adaptation and intercellular signaling. Overall, our findings indicate that the resting HUVEC secretome is thus composed of both conventionally and unconventionally secreted proteins, which contribute to coagulation, angiogenesis, and immune processes. Unexpectedly, we identified the adult hemoglobin subunit α-globin as a candidate UCPS cargo. The α-globin protein, encoded by the *HBA1* and *HBA2* genes, is expressed at high levels in erythroid precursors and binds β-globin to form adult hemoglobin (HbA, α2β2). Human and mouse endothelial cells are reported to express α-globin genes at much lower levels^37^ but secretion of the protein has not been recognized previously.

### Evaluation of UCPS prediction algorithms

Several computational tools have been developed to predict proteins secreted via UCPS, including SecretomeP^41^ and OutCyte^42^. SecretomeP was built using a neural network-based approach analyzing features within conventionally secreted proteins after N-terminal signal-peptide removal and posits that secreted proteins share physiochemical characteristics independent of the secretory pathway used. SecretomeP recommends a threshold score of 0.6 for classifying proteins as secreted. To assess its performance, we compared SecretomeP scores for proteins detected exclusively in the medium fraction (lacking signal peptides and resistant to brefeldin-A; **Fig. 2A**) with those displaying low M/L ratios and enriched in the lysate fraction.

**Figure 2.**
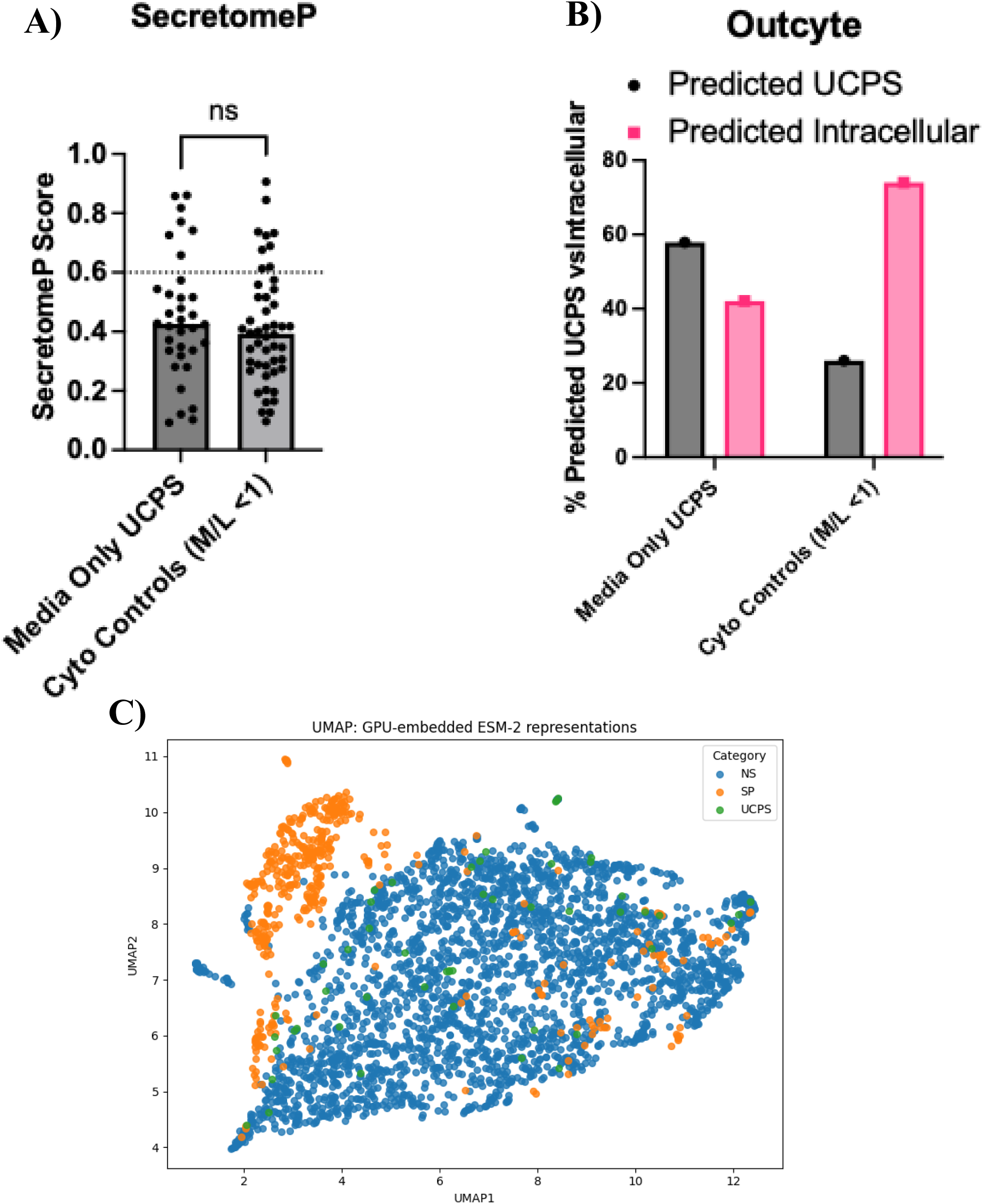
Comparison of computations tools for predicting unconventionally secreted cytoplasmic proteins. **A)** SecretomeP scores for proteins identified exclusively in the media fraction lacking N-terminal signal peptides (”Media Only UCPS”), and proteins enriched in the lysate fraction ( Cyto Controls with M/L <1, n = 3). **B)** Percentage of proteins classified as UCPS or intracellular by Outcyte in the Media Only UCPS and Cyto Control groups. **C)** UMAP projection of protein sequence embeddings generated by the ESM-2 language model for signal-peptide containing (SP), unconventionally secreted (UCPS), and non-secreted (NS) proteins.

SecretomeP failed to discriminate between these groups, with median scores of 0.45 and 0.41, respectively (**Fig. 2A**). OutCyte is a newer algorithm that leverages experimentally derived secretome data to identify conserved features among UCPS cargoes^42^. OutCyte showed improved discrimination relative to SecretomeP, classifying 58% of media-only proteins and 26% of lysate-enriched controls as unconventionally secreted (**Fig. 2B**). However, further analysis of OutCyte’s top predictive features, including relative arginine frequency, molecular weight, and the abundance of positively charged residues, revealed that these metrics did not significantly differ between UCPS cargoes and cytoplasmic control proteins (**Fig. S2)**. This suggests that while OutCyte outperforms SecretomeP, its underlying discriminative features may not robustly distinguish bona fide UCPS cargoes from cytoplasmic proteins. Both SecretomeP and OutCyte utilize traditional machine learning-based approaches that are constrained by predefined features, such as amino acid composition and hydrophobicity. In contrast, protein language models, such as the evolutionary-scale language model (ESM), generate sequence representations that implicitly encode structural, functional, and evolutionary properties, capturing the biological semantics learned from millions of protein sequences ^43^. We used ESM-2 (esm2_t33_650M_UR50D) to embed signal peptide-containing proteins (SP), UCPS cargoes, and non-secreted (NS) cytoplasmic proteins from the HUVEC dataset into numerical vectors ^43^. UMAP projection of these embeddings revealed a distinct clustering of SP proteins, while UCPS and NS proteins were highly intermixed (**Fig. 2C**). This indicates that UCPS cargoes do not exhibit global sequence features that clearly separate them from cytoplasmic proteins, underscoring the need for alternative approaches that incorporate context-dependent variables such as subcellular localization dynamics, post-translational modifications, and stress-induced signaling pathways to more accurately define UCPS cargoes.

### HUVEC secretome alterations induced by inflammatory stimuli

To examine the influence of inflammatory stress on the endothelial secretome, we treated HUVECs with 1 µg/mL lipopolysaccharide (LPS), a Gram-negative bacterial endotoxin commonly used to study inflammatory responses. Four hours after treatment with LPS, we observed minimal changes in the proteome and secretome of HUVECs, which is consistent with findings that TLR4 treatment of endothelial cells evokes slower but more sustained responses compared to innate immune cells ^44, 45^ (**Fig. S3A**). We therefore pretreated HUVECs for 20 hours with 1 µg/mL LPS in complete medium, followed by 4-hour incubation in serum-free conditions prior to collection of proteins for secretome analysis. Following this extended treatment, there was an increase in the cellular abundance of several classes of inflammatory proteins (**Fig. 3A**; green and purple datapoints indicate proteins containing and lacking signal peptides, respectively). This included activation of the non-canonical nuclear factor kappa B (NF-κB) inflammatory response, as indicated by upregulation of the nuclear factor NF-kappa-B p100 subunit (NFKB2) and its associated transcription factor RelB (RELB) ^46^, which has been shown to mediate endothelial inflammatory responses^47-49^. Several known NF-κB induced genes were also found to be up-regulated, including adhesion molecules (ICAM1, SELE, VCAM1) ^50, 51^ and chemokines (CCL2, CXCL6, CXCL8) ^52-55^, which aid in the recruitment and attachment of innate and adaptive immune cells to inflamed sites. Similarly, several proteins involved in the response to LPS triggered inflammation and the antimicrobial response exhibited increased secretion into the conditioned medium relative to untreated controls (**Fig. 3B**). This included the pro-inflammatory cytokine interleukin 6 (IL6), antimicrobial enzymes (CTSK, CTSS), chemoattractant proteins (CXCL1, CXCL2, CXLC6, CCL2, CCL20), alongside an increase in antigen presentation as indicated by an increase in HLA class I histocompatibility antigen, alpha chain E (HLA-E) ^56^ (**Supplemental File. 1)**. In contrast, only three proteins were found to have decreased abundance across lysate and medium fractions, underscoring the predominant effect of LPS in increasing the production of intracellular and secreted proteins involved in the inflammatory response.

**Figure 3.**
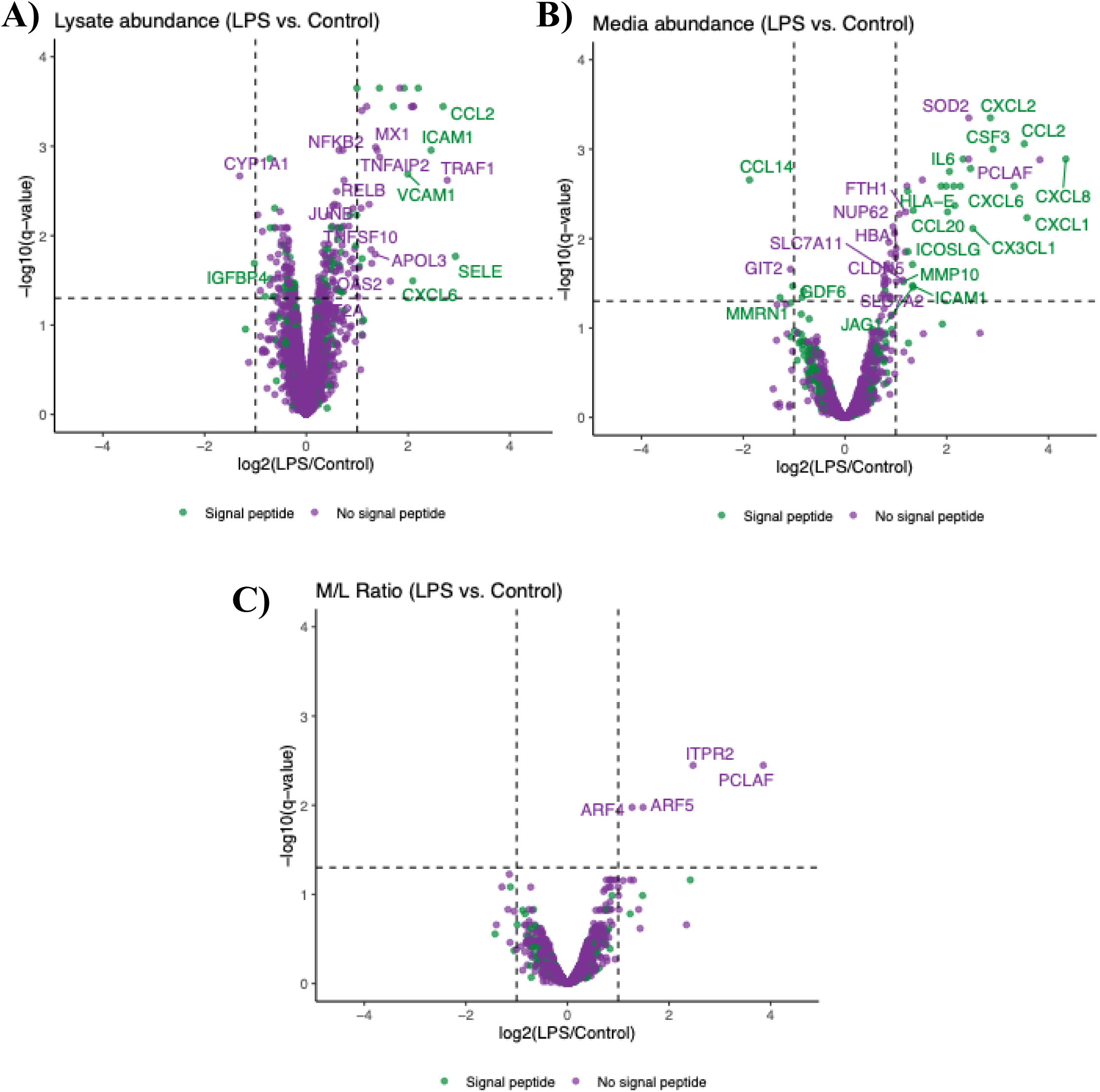
LPS treatment induces secretion of pro-inflammatory molecules from HUVEC. Volcano plots comparing protein abundance in the **(A)** lysate fraction, **(B)** media fraction, and **(C)** M/L ratios of HUVEC treated with 1ug.ml LPS versus control (n = 3 per group).

Of the 34 proteins with increased abundance in conditioned medium, the majority (80%, 28 proteins) contain signal peptides, indicating that they were conventionally secreted. Interestingly, most of these proteins were not detected in the lysate fraction (**Fig. 3A-B**). While this precludes our ability to calculate M/L ratios, it indicates that these proteins were secreted efficiently following synthesis. Furthermore, the proteins detected in both the medium and lysate fractions following LPS treatment (e.g., CXCL6, CCL2) had a higher abundance in both fractions, resulting in M/L ratios that were not significantly different from those of untreated controls (**Fig. 3C)**.

Two known unconventionally secreted proteins, superoxide dismutase 2 (SOD2) and ferritin heavy chain (FTH1) ^57-59^, were found to be more abundant in conditioned medium following LPS treatment. Interestingly, both FTH1 and SOD2 are known to regulate the redox status of the extracellular environment by either acting as an antioxidant or by catalyzing dismutation of the superoxide radical, respectively. FTH1 has been shown to be secreted from mouse oligodendrocytes and stem cell-derived blood-brain barrier endothelial cells^57, 59^. Our study indicates that FTH1 is also secreted by HUVECs, and that its secretion is stimulated by LPS treatment. SOD2 has previously been shown to be unconventionally secreted by macrophages^60^, and we found that its abundance in both the medium and lysate fractions are significantly increased following treatment with LPS.

ARF4, ARF4, NUP62, α-globin, and PCLAF are novel candidate UCPS cargoes identified in our dataset as they lack signal peptides and transmembrane domains but are not reported to be secreted (**Fig. 3 B-C**). Surprisingly, we found that α-globin, a component of adult hemoglobin, was also significantly more abundant in the medium following LPS treatment, indicating potential unconventional secretion of this protein. While α-globin is expressed at very high levels in erythrocytes, where it forms a tetrameric complex with β-globin to form adult hemoglobin (α2β2), low-level expression of globin proteins has also been described in non-erythroid cells, including endothelial cells^37, 61-63^.

Transcriptome analysis of HUVECs has previously identified NLRP3, PYCARD, and CASP1, components of the NLRP3 inflammasome. Thus, we next investigated whether the NLRP3 inflammasome contributes to the regulation of the endothelial secretome under inflammatory conditions. The NLRP3 inflammasome requires two signals for activation, with the first signal priming the synthesis of NLRP3 genes and cytokines (provided by LPS in our study), and the second stimulus, such as the ionophore nigericin, triggering assembly of the inflammasome complex and cytokine maturation. To test this hypothesis, we compared the secretome profiles of HUVECs treated with LPS alone, 1 hour treatment with nigericin alone, or LPS followed by nigericin. Despite LPS treatment, we did not detect NLRP3 protein or IL-1β in cell lysates, nor did we observe IL-1β secretion in any condition (**Supplementary File 1**). These findings suggest that HUVECs do not express sufficient levels of NLRP3 protein to support inflammasome assembly, despite transcriptional evidence of the component genes. However, treatment with nigericin – either alone or following LPS – resulted in increased secretion proteins via the conventional and UCPS pathways. Among these, six proteins (AHSG, APOC3, CTSL, α-globin, HBD, and ITIH4) displayed M/L ratios greater > 1 following combined LPS/nigericin treatment (**Fig. S3 A**). However, four of these six proteins (AHSG, CTSL, α-globin, and HBD) were also elevated following nigericin treatment alone (**Fig. S3 B**), indicating that their secretion was primarily driven by nigericin rather than synergistic inflammasome activation. These findings suggest that brief exposure to nigericin promotes release of a subset of unconventionally secreted proteins in HUVECs through a mechanism independent of NLRP3. This likely reflects ionophore-induced cellular stress or membrane perturbation rather than canonical inflammasome activation.

### Validation of the novel UCPS cargo, α-globin

Mass spectrometry analysis identified α-globin as a UCPS cargo released from resting HUVECs, with secretion being potentiated following treatment with LPS and nigericin. We thus sought to confirm the presence of α-globin in HUVEC conditioned medium using immunoblotting with a previously characterized polyclonal anti-α-globin antibody^62^. Consistent with the mass spectrometry analysis, α-globin was predominantly detected in the conditioned medium, and its concentration increased following LPS and LPS/Nigericin treatment (**Fig. 4A lanes 1-3**). Treatment of cells with brefeldin-A did not impair α-globin secretion (**Fig. 4A lane 4**). As a control, we analyzed the secretion of interleukin-8 (CXCL8), a LPS-sensitive, conventionally secreted protein identified in our secretome analysis. Interleukin-8 synthesis and secretion were markedly increased following LPS treatment, and secretion was reduced following treatment with BFA (**Fig. 4A**). These findings validate that α-globin is unconventionally secreted by HUVECs, and its secretion is enhanced upon LPS and nigericin stimulation.

**Figure 4.**
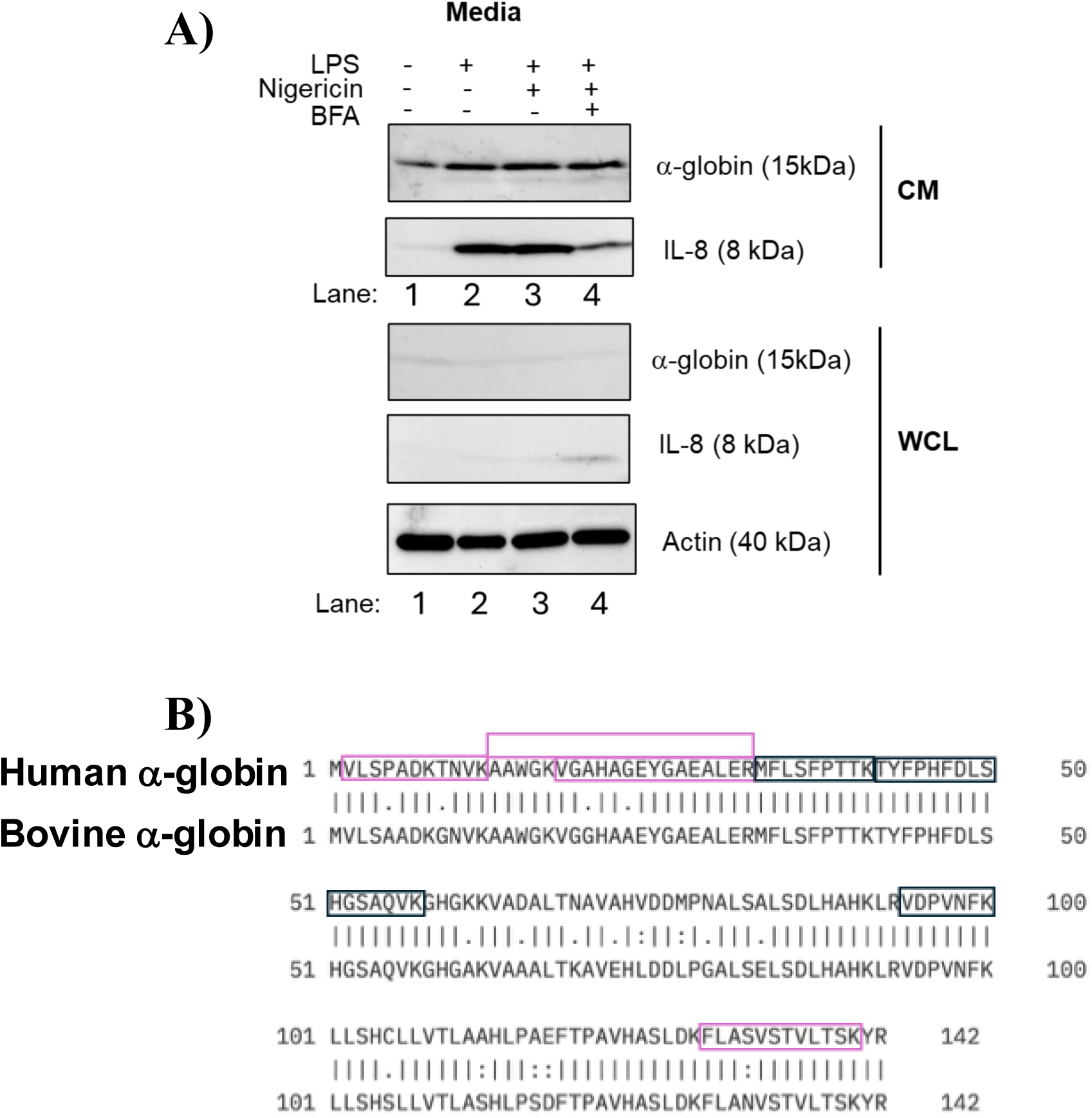
a-globin is unconventionally secreted by HUVECs. **A)** Immunoblotting analysis of a-globin and IL-8 in HUVEC whole cell lysates (WCL) and conditioned media (CM) following treatment with 1 ug.ml LPS, 1 uM Nigericin, and 1 ug.ml brefeldin-A (BFA). **B)** Aligned human and bovine α-globin sequences, with peptides identified in our study boxed, and human-specific peptides highlighted with red boxes.

To address the possibility of contamination by bovine α-globin, as HUVECs are cultured in fetal bovine serum prior to experiments conducted in serum-free medium, we analyzed the α-globin peptides identified in our dataset. Human and bovine α-globin share 88% sequence identity; however, across three independent datasets, we identified five human-specific α-globin peptides in the conditioned medium (**Fig. 4B**; human-specific peptides are boxed in red). Of the five human-specific peptides detected, four were significantly more enriched in conditioned medium compared to cell lysates, confirming the secretion of human α-globin by HUVECs (**Fig S4**). Considered together, these data support unconventional secretion of α-globin from HUVECs.

## Discussion

In this study, we performed quantitative proteomic profiling of the cellular proteome and secretome of primary human umbilical vein endothelial cells (HUVECs) under basal conditions and following stimulation with LPS and nigericin. By comparing protein abundance in cell lysates and conditioned medium, we identified proteins enriched in the medium relative to the lysate, enabling discrimination between actively secreted proteins and intracellular contaminants ^26, 27^, and treatment with BFA, an inhibitor of ER-Golgi trafficking, confirmed BFA-resistant, unconventional secretion of cytoplasmic proteins. Contrary to the assumption that inflammatory stimuli broadly enhance unconventional protein secretion (UCPS) ^16, 21^, our data revealed a more selective pattern. While numerous proteins increased in abundance in conditioned medium following LPS stimulation, most contained signal peptides, consistent with secretion via the classical ER-Golgi pathway. Only a small subset of signal-peptide-lacking proteins, including superoxide dismutase 2 (SOD2) and α-globin were enriched in the secretome following stimulation. This suggests that LPS does not globally activate UCPS pathways in endothelial cells but may instead promote selective secretion of specific cargoes.

This study utilized a single concentration and timepoint for LPS and nigericin treatment, representing a snapshot of the endothelial secretory response. Future studies could apply dose-response and time-course analyses to assess how the relative contribution of conventional and unconventional secretory pathways is shaped by the strength and duration of inflammatory stimuli. In addition, distinct pathogen-associated molecular patterns or danger-associated molecular patterns sensed by specific pathogen recognition receptors, may trigger divergent secretion pathways, further highlighting the need for stimulus-specific profiling. For example, endothelial cells have been documented to express RNA sensing PRR’s TLR3 and RIG-I, and several isoforms of the cytosolic NLR family members^64, 65^. Furthermore, endothelial cells exhibit transcriptional and functional heterogeneity across vascular beds^66^. Single-cell RNA-sequencing and bulk transcriptomic studies have demonstrated distinct gene expression signatures that underlie organ-specific vascular phenotypes^67-69^. Applying secretome analyses to endothelial cells isolated from different tissues may uncover previously unrecognized secreted factors that contribute to tissue-specific functions, including leukocyte recruitment, vascular permeability, and angiocrine signaling.

Previous secretome studies in immortalized endothelial lines such as EA.hy926^70, 71^ cells have reported the secretion of α-globin in response to LPS (see supplementary table in Songjang et al. ^70^). Our study extends these observations to primary HUVECs. By directly comparing protein abundance in medium and lysate fractions, combined with immunoblotting analysis, we confirmed that α-globin is robustly secreted following LPS stimulation, suggesting that its release may be a conserved feature of the endothelial inflammatory response. Red blood cell precursors express very high levels of α-globin, which combines with β-globin to form HbA (α2β2), the blood oxygen carrier. Human and mouse endothelial cells also express α-globin, albeit at much lower levels ^37,63^. Although these levels are too low to facilitate oxygen transport, α-globin can also function as an enzyme through electron transport mediated by its associated heme moiety. In this capacity, α-globin can generate reactive oxygen species from heme-bound oxygen or participate in additional reduction and oxidation (redox) reactions through interactions with other heme ligands, including nitric oxide (NO), peroxides, and nitrite (NO2^−^). In endothelial cells, α-globin located in the myoendothelial junction is thought to enhance vascular contractility by degrading the vasodilator nitric oxide ^62, 72, 73^. Analogous to other unconventionally secreted redox-regulating proteins^74^, including superoxide dismutase 1^75^, thioredoxin^76^, and thioredoxin reductase^77^, endothelial cell-secreted α-globin may contribute to modulation of the local extracellular redox environment, particularly at sites of inflammation. Given the critical role of extracellular redox balance in regulating endothelial barrier function^78, 79^ and leukocyte adhesion^80^, α-globin secretion may represent a previously unappreciated mechanism of inflammatory control.

While the structural features that dictate entry into conventional secretory pathways are well described, there are no known shared features among UCPS cargoes that direct them toward secretion. Computational tools such as SecretomeP^41^ and OutCyte^42^ were unable to reliably distinguish between UCPS cargoes and control non-secreted cytoplasmic proteins in our dataset. These tools were built by analyzing primary amino acid sequences and classical machine learning frameworks, and do not incorporate post-translational modifications or protein secondary/tertiary structural features that may be critical for entry into unconventional secretion pathways. The development of advanced models based on deep learning and structural prediction, such as those leveraging protein language models or AlphaFold embeddings, may provide more accurate predictions of unconventional secretion, particularly when integrated with proteomic datasets. Our study provides a systematic analysis of the HUVEC secretome under resting and inflammatory conditions, and reveals selective, rather than global, activation of unconventional protein secretion in response to LPS. The robust secretion of α-globin and other non-canonical cargoes highlights the need for deeper mechanistic understanding of UCPS in endothelial cells. These findings lay the groundwork for future efforts to dissect tissue-specific endothelial secretomes and to develop improved computational tools for predicting secretion via unconventional secretory pathways.

## Abbreviations

TLR4: Toll-like receptor 4
ER: Endoplasmic Reticulum
UCPS: Unconventional Cytoplasmic Protein Secretion
HUVEC: Human Umbilical Vein Endothelial Cells
HBA1: Hemoglobin Subunit Alpha
TMT: Tandem Mass Tag
BFA: Brefeldin A
LPS: Lipopolysaccharide.

## Acknowledgments

This work was funded by National Institutes of Health (NIH), National Institute of General Medical Sciences grant R00GM141268 (P.S.A.), and R01-GM-094231 and U24-CA271037 (A.I.N). National Heart, Lung, and Blood Institute grants R01HL165798, R01HL156647, and U01HL163983 (M.J.W.), The Assisi Foundation (M.J.W.), and the American Lebanese Syrian Associated Charities (M.J.W). This work was supported by the Canadian Institutes of Health Research (PJT-168987, to C.A.K.). We acknowledge the University of Michigan’s Proteomics Resource Facility (RRID: SCR_026723) for providing assistance with proteomic data acquisition.

---

**Supplemental Figure 1.**
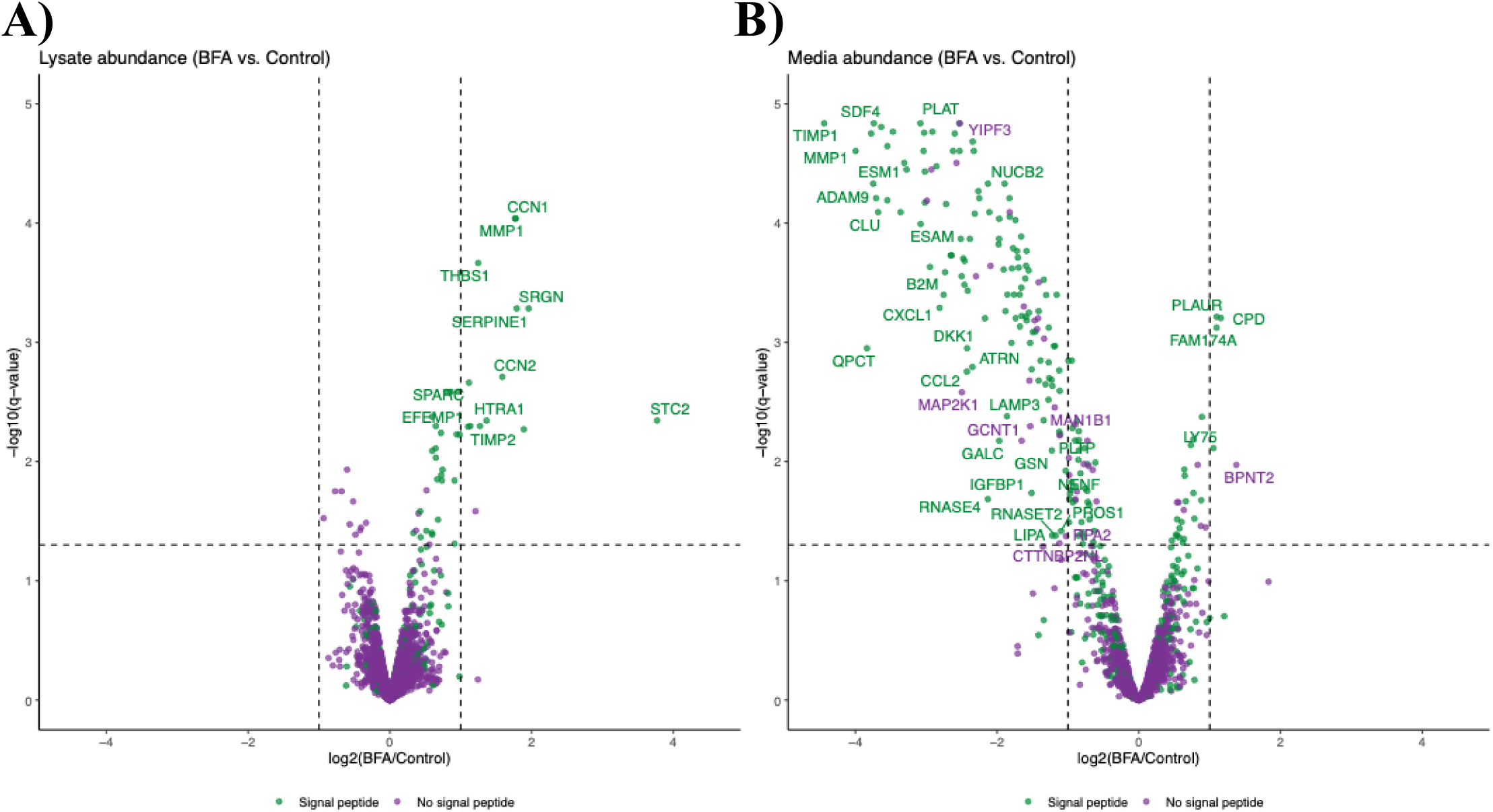
Brefeldin-A treatment causes accumulation of signal-peptide containing proteins in the cell lysate fraction, and their depletion from the conditioned media. Volcano plots comparing protein abundance in the cell lysate (n = 3 per group) **(A)** and conditioned media (n = 3 per group) **(B)** fractions following treatment with brefeldin-A versus control.

**Supplemental Figure 2.**
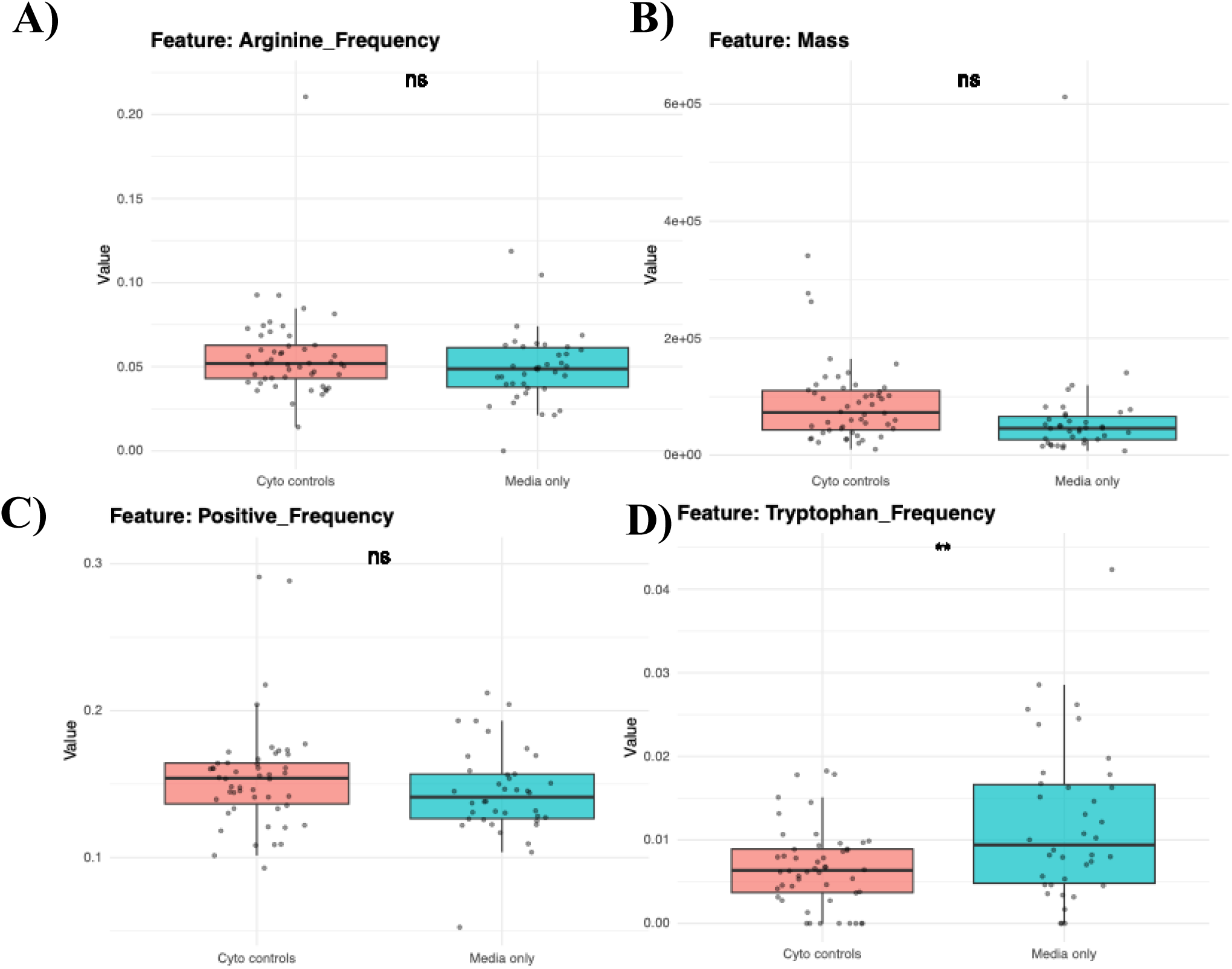
Key features identified by Outcyte do not significantly differ between putative UCPS cargoes and control cytoplasmic proteins. Analysis of A) arginine frequency, B) mass, C) positive amino acid frequency, and D) tryptophan frequency in a control group of proteins with low M/L ratios (<1; “Cyto control”), and proteins detected only in the conditioned media (Media Only).

**Figure 3 supplement 1.**
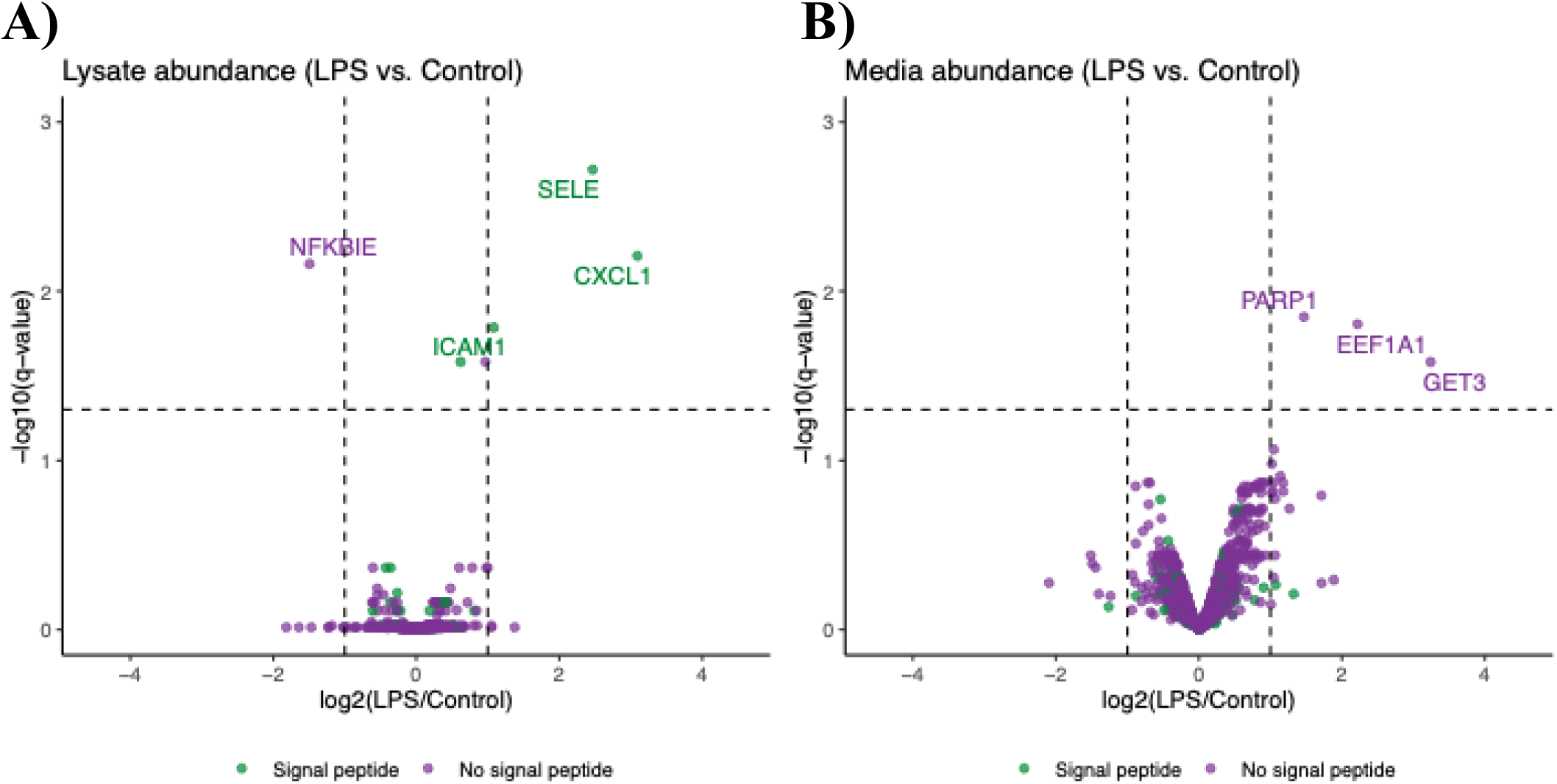
4 h treatment with 1ug/ml LPS has minor impact on A) HUVEC lysate and B) secretome (n = 3 per group).

**Figure 3 supplement 2.**
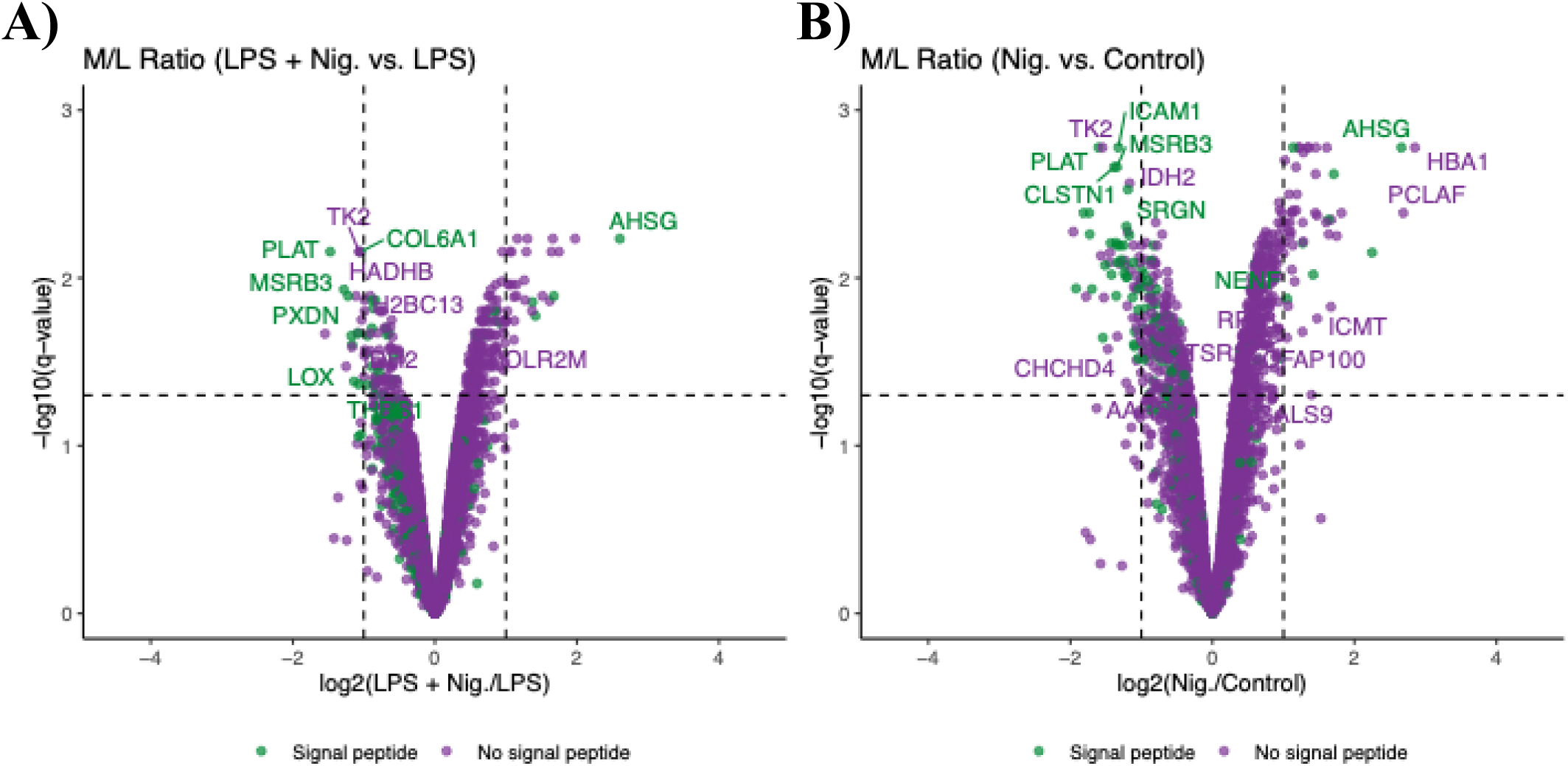
Treatment of HUVEC with nigericin predominantly increases secretion of signal-peptide lacking proteins. **A)** Volcano plots comparing M/L ratios of HUVEC treated with 1 uM nigericin compared to control untreated cells, and treatment with **B)** LPS following an LPS priming phase compared to LPS treatment alone (n = 3 per group).

**Figure 4 supplement.**
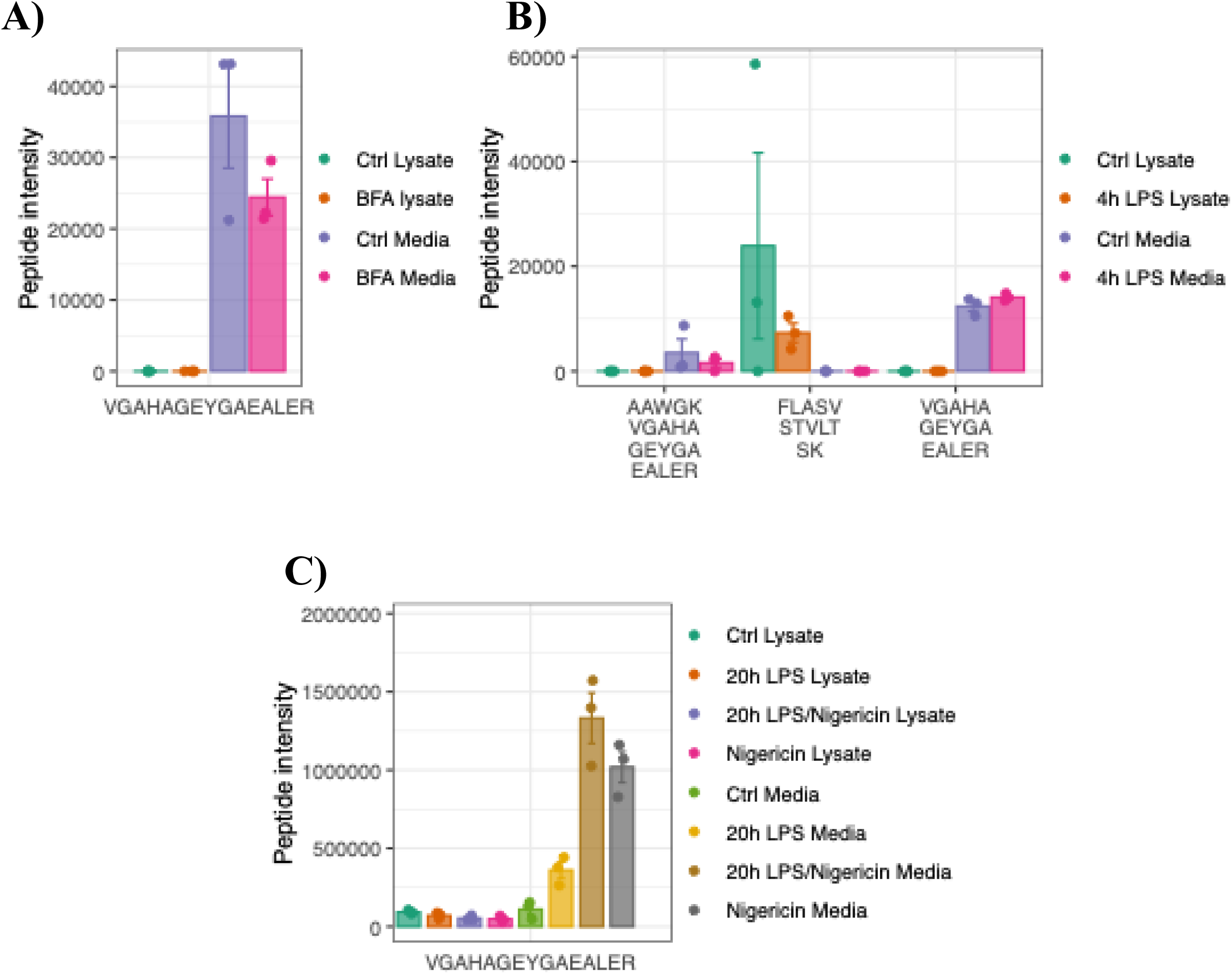
Human-specific α-globin peptides are enriched in the HUVEC conditioned media. **A-C)** Relative abundance of human-specific α-globin peptides identified in HUVEC cell conditioned media and lysate in three independent datasets.

## References

1. Michiels C. Endothelial cell functions. Journal of cellular physiology. 2003;196(3):430–43. doi: 10.1002/jcp.10333. PubMed PMID: 12891700.

2. Muller WA. Leukocyte-endothelial cell interactions in the inflammatory response. Lab Invest. 2002;82(5):521–33. doi: 10.1038/labinvest.3780446. PubMed PMID: 12003992.

3. Amersfoort J, Eelen G, Carmeliet P. Immunomodulation by endothelial cells - partnering up with the immune system? Nat Rev Immunol. 2022;22(9):576–88. Epub 20220314. doi: 10.1038/s41577-022-00694-4. PubMed PMID: 35288707; PMCID: PMC8920067.

4. Zhou H, Andonegui G, Wong CH, Kubes P. Role of endothelial TLR4 for neutrophil recruitment into central nervous system microvessels in systemic inflammation. Journal of immunology. 2009;183(8):5244–50. Epub 20090928. doi: 10.4049/jimmunol.0901309. PubMed PMID: 19786543.

5. Wang W, Deng M, Liu X, Ai W, Tang Q, Hu J. TLR4 activation induces nontolerant inflammatory response in endothelial cells. Inflammation. 2011;34(6):509–18. doi: 10.1007/s10753-010-9258-4. PubMed PMID: 20878353.

6. Yazji I, Sodhi CP, Lee EK, Good M, Egan CE, Afrazi A, Neal MD, Jia H, Lin J, Ma C, Branca MF, Prindle T, Richardson WM, Ozolek J, Billiar TR, Binion DG, Gladwin MT, Hackam DJ. Endothelial TLR4 activation impairs intestinal microcirculatory perfusion in necrotizing enterocolitis via eNOS-NO-nitrite signaling. Proceedings of the National Academy of Sciences of the United States of America. 2013;110(23):9451–6. Epub 20130506. doi: 10.1073/pnas.1219997110. PubMed PMID: 23650378; PMCID: PMC3677476.

7. Andonegui G, Zhou H, Bullard D, Kelly MM, Mullaly SC, McDonald B, Long EM, Robbins SM, Kubes P. Mice that exclusively express TLR4 on endothelial cells can efficiently clear a lethal systemic Gram-negative bacterial infection. Journal of Clinical Investigation. 2009;119(7):1921–30. doi: 10.1172/Jci36411. PubMed PMID: WOS:000267694300021.

8. Salvador B, Arranz A, Francisco S, Cordoba L, Punzon C, Llamas MA, Fresno M. Modulation of endothelial function by Toll like receptors. Pharmacol Res. 2016;108:46–56. Epub 20160409. doi: 10.1016/j.phrs.2016.03.038. PubMed PMID: 27073018.

9. Nagyoszi P, Nyul-Toth A, Fazakas C, Wilhelm I, Kozma M, Molnar J, Hasko J, Krizbai IA. Regulation of NOD-like receptors and inflammasome activation in cerebral endothelial cells. Journal of neurochemistry. 2015;135(3):551–64. Epub 20150903. doi: 10.1111/jnc.13197. PubMed PMID: 26083549.

10. Bleda S, de Haro J, Varela C, Esparza L, Ferruelo A, Acin F. NLRP1 inflammasome, and not NLRP3, is the key in the shift to proinflammatory state on endothelial cells in peripheral arterial disease. Int J Cardiol. 2014;172(2):e282–4. Epub 20140109. doi: 10.1016/j.ijcard.2013.12.201. PubMed PMID: 24439873.

11. Bai B, Yang Y, Wang Q, Li M, Tian C, Liu Y, Aung LHH, Li PF, Yu T, Chu XM. NLRP3 inflammasome in endothelial dysfunction. Cell death & disease. 2020;11(9):776. Epub 20200918. doi: 10.1038/s41419-020-02985-x. PubMed PMID: 32948742; PMCID: PMC7501262.

12. Carman CV, Martinelli R. T Lymphocyte-Endothelial Interactions: Emerging Understanding of Trafficking and Antigen-Specific Immunity. Frontiers in immunology. 2015;6:603. Epub 20151124. doi: 10.3389/fimmu.2015.00603. PubMed PMID: 26635815; PMCID: PMC4657048.

13. Mai J, Virtue A, Shen J, Wang H, Yang XF. An evolving new paradigm: endothelial cells--conditional innate immune cells. J Hematol Oncol. 2013;6:61. Epub 20130822. doi: 10.1186/1756-8722-6-61. PubMed PMID: 23965413; PMCID: PMC3765446.

14. Araujo TLS, Zeidler JD, Oliveira PVS, Dias MH, Armelin HA, Laurindo FRM. Protein disulfide isomerase externalization in endothelial cells follows classical and unconventional routes. Free Radic Biol Med. 2017;103:199–208. Epub 20161227. doi: 10.1016/j.freeradbiomed.2016.12.021. PubMed PMID: 28034831.

15. Pallet N, Sirois I, Bell C, Hanafi LA, Hamelin K, Dieude M, Rondeau C, Thibault P, Desjardins M, Hebert MJ. A comprehensive characterization of membrane vesicles released by autophagic human endothelial cells. Proteomics. 2013;13(7):1108–20. doi: 10.1002/pmic.201200531. PubMed PMID: 23436686.

16. Rabouille C. Pathways of Unconventional Protein Secretion. Trends in cell biology. 2017;27(3):230–40. doi: 10.1016/j.tcb.2016.11.007. PubMed PMID: 27989656.

17. Zanetti G, Pahuja KB, Studer S, Shim S, Schekman R. COPII and the regulation of protein sorting in mammals. Nature cell biology. 2011;14(1):20–8. doi: 10.1038/ncb2390. PubMed PMID: 22193160.

18. Schekman RW. Retrospective: George E. Palade (1912-2008). Science. 2008;322(5902):695. doi: 10.1126/science.1167174. PubMed PMID: 18974342.

19. Nightingale T, Cutler D. The secretion of von Willebrand factor from endothelial cells; an increasingly complicated story. Journal of thrombosis and haemostasis : JTH. 2013;11 Suppl 1(Suppl 1):192-201. doi: 10.1111/jth.12225. PubMed PMID: 23809123; PMCID: PMC4255685.

20. Lenting PJ, Christophe OD, Denis CV. von Willebrand factor biosynthesis, secretion, and clearance: connecting the far ends. Blood. 2015;125(13):2019–28. Epub 20150223. doi: 10.1182/blood-2014-06-528406. PubMed PMID: 25712991.

21. Kim J, Gee HY, Lee MG. Unconventional protein secretion - new insights into the pathogenesis and therapeutic targets of human diseases. Journal of cell science. 2018;131(12). doi: 10.1242/jcs.213686. PubMed PMID: 29941450.

22. Neel E, Chiritoiu-Butnaru M, Fargues W, Denus M, Colladant M, Filaquier A, Stewart SE, Lehmann S, Zurzolo C, Rubinsztein DC, Marin P, Parmentier ML, Villeneuve J. The endolysosomal system in conventional and unconventional protein secretion. The Journal of cell biology. 2024;223(9). Epub 20240812. doi: 10.1083/jcb.202404152. PubMed PMID: 39133205; PMCID: PMC11318669.

23. Broz P. Unconventional protein secretion by gasdermin pores. Semin Immunol. 2023;69:101811. Epub 20230718. doi: 10.1016/j.smim.2023.101811. PubMed PMID: 37473560.

24. Abbineni PS, Tang VT, da Veiga Leprevost F, Basrur V, Xiang J, Nesvizhskii AI, Ginsburg D. Identification of secreted proteins by comparison of protein abundance in conditioned media and cell lysates. Anal Biochem. 2022;655:114846. Epub 20220813. doi: 10.1016/j.ab.2022.114846. PubMed PMID: 35973625; PMCID: PMC9756135.

25. Schira-Heinen J, Grube L, Waldera-Lupa DM, Baberg F, Langini M, Etemad-Parishanzadeh O, Poschmann G, Stuhler K. Pitfalls and opportunities in the characterization of unconventionally secreted proteins by secretome analysis. Biochimica et biophysica acta Proteins and proteomics. 2019;1867(12):140237. doi: 10.1016/j.bbapap.2019.06.004. PubMed PMID: 31202002.

26. Tang VT, Abbineni PS, Veiga Leprevost FD, Basrur V, Khoriaty R, Emmer BT, Nesvizhskii AI, Ginsburg D. Identification of LMAN1- and SURF4-Dependent Secretory Cargoes. Journal of proteome research. 2023;22(11):3439–46. Epub 20231016. doi: 10.1021/acs.jproteome.3c00259. PubMed PMID: 37844105; PMCID: PMC10629478.

27. Abbineni PS, Tang VT, da Veiga Leprevost F, Basrur V, Xiang J, Nesvizhskii AI, Ginsburg D. Identification of secreted proteins by comparison of protein abundance in conditioned media and cell lysates. bioRxiv. 2022:2022.06.16.496407. doi: 10.1101/2022.06.16.496407.

28. Poschmann G, Brenig K, Lenz T, Stuhler K. Comparative Secretomics Gives Access to High Confident Secretome Data: Evaluation of Different Methods for the Determination of Bona Fide Secreted Proteins. Proteomics. 2021;21(2):e2000178. doi: 10.1002/pmic.202000178. PubMed PMID: 33015975.

29. Grube L, Dellen R, Kruse F, Schwender H, Stuhler K, Poschmann G. Mining the Secretome of C2C12 Muscle Cells: Data Dependent Experimental Approach To Analyze Protein Secretion Using Label-Free Quantification and Peptide Based Analysis. Journal of proteome research. 2018;17(2):879–90. doi: 10.1021/acs.jproteome.7b00684. PubMed PMID: 29322779.

30. Ritchie ME, Phipson B, Wu D, Hu Y, Law CW, Shi W, Smyth GK. limma powers differential expression analyses for RNA-sequencing and microarray studies. Nucleic acids research. 2015;43(7):e47. Epub 20150120. doi: 10.1093/nar/gkv007. PubMed PMID: 25605792; PMCID: PMC4402510.

31. Ryckman C, Vandal K, Rouleau P, Talbot M, Tessier PA. Proinflammatory activities of S100: proteins S100A8, S100A9, and S100A8/A9 induce neutrophil chemotaxis and adhesion. Journal of immunology. 2003;170(6):3233-42. doi: 10.4049/jimmunol.170.6.3233. PubMed PMID: 12626582.

32. Burgener SS, Leborgne NGF, Snipas SJ, Salvesen GS, Bird PI, Benarafa C. Cathepsin G Inhibition by Serpinb1 and Serpinb6 Prevents Programmed Necrosis in Neutrophils and Monocytes and Reduces GSDMD-Driven Inflammation. Cell reports. 2019;27(12):3646–56 e5. doi: 10.1016/j.celrep.2019.05.065. PubMed PMID: 31216481; PMCID: PMC7350907.

33. Sanrattana W, Maas C, de Maat S. SERPINs-From Trap to Treatment. Front Med (Lausanne). 2019;6:25. Epub 20190212. doi: 10.3389/fmed.2019.00025. PubMed PMID: 30809526; PMCID: PMC6379291.

34. Baumann M, Pham CT, Benarafa C. SerpinB1 is critical for neutrophil survival through cell-autonomous inhibition of cathepsin G. Blood. 2013;121(19):3900–7, S1-6. Epub 20130326. doi: 10.1182/blood-2012-09-455022. PubMed PMID: 23532733; PMCID: PMC3650706.

35. El Ouaamari A, Dirice E, Gedeon N, Hu J, Zhou JY, Shirakawa J, Hou L, Goodman J, Karampelias C, Qiang G, Boucher J, Martinez R, Gritsenko MA, De Jesus DF, Kahraman S, Bhatt S, Smith RD, Beer HD, Jungtrakoon P, Gong Y, Goldfine AB, Liew CW, Doria A, Andersson O, Qian WJ, Remold-O’Donnell E, Kulkarni RN. SerpinB1 Promotes Pancreatic beta Cell Proliferation. Cell Metab. 2016;23(1):194–205. Epub 20151215. doi: 10.1016/j.cmet.2015.12.001. PubMed PMID: 26701651; PMCID: PMC4715773.

36. Liu JZ, Hu YL, Feng Y, Jiang Y, Guo YB, Liu YF, Chen X, Yang JL, Chen YY, Mao QS, Xue WJ. BDH2 triggers ROS-induced cell death and autophagy by promoting Nrf2 ubiquitination in gastric cancer. J Exp Clin Cancer Res. 2020;39(1):123. Epub 20200630. doi: 10.1186/s13046-020-01620-z. PubMed PMID: 32605589; PMCID: PMC7325376.

37. Abbineni PS, Baid S, Weiss MJ. A moonlighting job for alpha-globin in blood vessels. Blood. 2024;144(8):834–44. doi: 10.1182/blood.2023022192. PubMed PMID: 38848504; PMCID: PMC11830976.

38. Riise R, Odqvist L, Mattsson J, Monkley S, Abdillahi SM, Tyrchan C, Muthas D, Yrlid LF. Bleomycin hydrolase regulates the release of chemokines important for inflammation and wound healing by keratinocytes. Scientific reports. 2019;9(1):20407. Epub 20191231. doi: 10.1038/s41598-019-56667-6. PubMed PMID: 31892708; PMCID: PMC6938525.

39. Wei N, Serino G, Deng XW. The COP9 signalosome: more than a protease. Trends Biochem Sci. 2008;33(12):592–600. Epub 20081014. doi: 10.1016/j.tibs.2008.09.004. PubMed PMID: 18926707.

40. Kuruba B, Starks N, Josten MR, Naveh O, Wayman G, Mikhaylova M, Kostyukova AS. Effects of Tropomodulin 2 on Dendritic Spine Reorganization and Dynamics. Biomolecules. 2023;13(8). Epub 20230811. doi: 10.3390/biom13081237. PubMed PMID: 37627302; PMCID: PMC10515316.

41. Bendtsen JD, Jensen LJ, Blom N, Von Heijne G, Brunak S. Feature-based prediction of non-classical and leaderless protein secretion. Protein engineering, design & selection : PEDS. 2004;17(4):349–56. doi: 10.1093/protein/gzh037. PubMed PMID: 15115854.

42. Zhao L, Poschmann G, Waldera-Lupa D, Rafiee N, Kollmann M, Stuhler K. OutCyte: a novel tool for predicting unconventional protein secretion. Scientific reports. 2019;9(1):19448. Epub 20191219. doi: 10.1038/s41598-019-55351-z. PubMed PMID: 31857603; PMCID: PMC6923414.

43. Lin Z, Akin H, Rao R, Hie B, Zhu Z, Lu W, Smetanin N, Verkuil R, Kabeli O, Shmueli Y, Dos Santos Costa A, Fazel-Zarandi M, Sercu T, Candido S, Rives A. Evolutionary-scale prediction of atomic-level protein structure with a language model. Science. 2023;379(6637):1123–30. Epub 20230316. doi: 10.1126/science.ade2574. PubMed PMID: 36927031.

44. Sawa Y, Ueki T, Hata M, Iwasawa K, Tsuruga E, Kojima H, Ishikawa H, Yoshida S. LPS-induced IL-6, IL-8, VCAM-1, and ICAM-1 expression in human lymphatic endothelium. J Histochem Cytochem. 2008;56(2):97-109. Epub 20071015. doi: 10.1369/jhc.7A7299.2007. PubMed PMID: 17938282; PMCID: PMC2324174.

45. Li M, van Esch B, Henricks PAJ, Garssen J, Folkerts G. Time and Concentration Dependent Effects of Short Chain Fatty Acids on Lipopolysaccharide- or Tumor Necrosis Factor alpha-Induced Endothelial Activation. Front Pharmacol. 2018;9:233. Epub 20180319. doi: 10.3389/fphar.2018.00233. PubMed PMID: 29615908; PMCID: PMC5867315.

46. Sun SC. The non-canonical NF-kappaB pathway in immunity and inflammation. Nat Rev Immunol. 2017;17(9):545–58. Epub 20170605. doi: 10.1038/nri.2017.52. PubMed PMID: 28580957; PMCID: PMC5753586.

47. Fan X, Li Q, Wang Y, Zhang DM, Zhou J, Chen Q, Sheng L, Passerini AG, Sun C. Non-canonical NF-kappaB contributes to endothelial pyroptosis and atherogenesis dependent on IRF-1. Transl Res. 2023;255:1–13. Epub 20221113. doi: 10.1016/j.trsl.2022.11.001. PubMed PMID: 36384204.

48. Jane-wit D, Surovtseva YV, Qin L, Li G, Liu R, Clark P, Manes TD, Wang C, Kashgarian M, Kirkiles-Smith NC, Tellides G, Pober JS. Complement membrane attack complexes activate noncanonical NF-kappaB by forming an Akt+ NIK+ signalosome on Rab5+ endosomes. Proceedings of the National Academy of Sciences of the United States of America. 2015;112(31):9686–91. Epub 20150720. doi: 10.1073/pnas.1503535112. PubMed PMID: 26195760; PMCID: PMC4534258.

49. Maracle CX, Agca R, Helder B, Meeuwsen JAL, Niessen HWM, Biessen EAL, de Winther MPJ, de Jager SCA, Nurmohamed MT, Tas SW. Noncanonical NF-kappaB signaling in microvessels of atherosclerotic lesions is associated with inflammation, atheromatous plaque morphology and myocardial infarction. Atherosclerosis. 2018;270:33–41. Epub 20180131. doi: 10.1016/j.atherosclerosis.2018.01.032. PubMed PMID: 29407886.

50. Rajan S, Ye J, Bai S, Huang F, Guo YL. NF-kappaB, but not p38 MAP kinase, is required for TNF-alpha-induced expression of cell adhesion molecules in endothelial cells. J Cell Biochem. 2008;105(2):477–86. doi: 10.1002/jcb.21845. PubMed PMID: 18613029; PMCID: PMC4422387.

51. Collins T, Read MA, Neish AS, Whitley MZ, Thanos D, Maniatis T. Transcriptional regulation of endothelial cell adhesion molecules: NF-kappa B and cytokine-inducible enhancers. FASEB J. 1995;9(10):899–909. PubMed PMID: 7542214.

52. Mukaida N, Okamoto S, Ishikawa Y, Matsushima K. Molecular mechanism of interleukin-8 gene expression. J Leukoc Biol. 1994;56(5):554–8. PubMed PMID: 7525815.

53. Kunsch C, Rosen CA. NF-kappa B subunit-specific regulation of the interleukin-8 promoter. Molecular and cellular biology. 1993;13(10):6137–46. doi: 10.1128/mcb.13.10.6137-6146.1993. PubMed PMID: 8413215; PMCID: PMC364673.

54. Wolter S, Doerrie A, Weber A, Schneider H, Hoffmann E, von der Ohe J, Bakiri L, Wagner EF, Resch K, Kracht M. c-Jun controls histone modifications, NF-kappaB recruitment, and RNA polymerase II function to activate the ccl2 gene. Molecular and cellular biology. 2008;28(13):4407-23. Epub 20080428. doi: 10.1128/MCB.00535-07. PubMed PMID: 18443042; PMCID: PMC2447141.

55. Richmond A. Nf-kappa B, chemokine gene transcription and tumour growth. Nat Rev Immunol. 2002;2(9):664–74. doi: 10.1038/nri887. PubMed PMID: 12209135; PMCID: PMC2668257.

56. Takao S, Ishikawa T, Yamashita K, Uchiyama T. The rapid induction of HLA-E is essential for the survival of antigen-activated naive CD4 T cells from attack by NK cells. Journal of immunology. 2010;185(10):6031–40. Epub 20101015. doi: 10.4049/jimmunol.1000176. PubMed PMID: 20952676.

57. Mukherjee C, Kling T, Russo B, Miebach K, Kess E, Schifferer M, Pedro LD, Weikert U, Fard MK, Kannaiyan N, Rossner M, Aicher ML, Goebbels S, Nave KA, Kramer-Albers EM, Schneider A, Simons M. Oligodendrocytes Provide Antioxidant Defense Function for Neurons by Secreting Ferritin Heavy Chain. Cell Metab. 2020;32(2):259–72 e10. Epub 20200611. doi: 10.1016/j.cmet.2020.05.019. PubMed PMID: 32531201; PMCID: PMC7116799.

58. Kimura T, Jia J, Kumar S, Choi SW, Gu Y, Mudd M, Dupont N, Jiang S, Peters R, Farzam F, Jain A, Lidke KA, Adams CM, Johansen T, Deretic V. Dedicated SNAREs and specialized TRIM cargo receptors mediate secretory autophagy. The EMBO journal. 2017;36(1):42–60. doi: 10.15252/embj.201695081. PubMed PMID: 27932448; PMCID: 5210154.

59. Palsa K, Baringer SL, Shenoy G, Spiegelman VS, Simpson IA, Connor JR. Exosomes are involved in iron transport from human blood-brain barrier endothelial cells and are modified by endothelial cell iron status. The Journal of biological chemistry. 2023;299(2):102868. Epub 20230103. doi: 10.1016/j.jbc.2022.102868. PubMed PMID: 36603765; PMCID: PMC9929479.

60. Chen J, Xu X, Shao Y, Bian X, Li R, Zhang Y, Xiao Y, Lu M, Jiang Q, Zeng Y, Yan F, Ye J, Li Z. AKT2 deficiency alleviates doxorubicin-induced cardiac injury via alleviating oxidative stress in cardiomyocytes. The international journal of biochemistry & cell biology. 2024;169:106539. Epub 20240128. doi: 10.1016/j.biocel.2024.106539. PubMed PMID: 38290690.

61. Keller TCSt, Lechauve C, Keller AS, Broseghini-Filho GB, Butcher JT, Askew Page HR, Islam A, Tan ZY, DeLalio LJ, Brooks S, Sharma P, Hong K, Xu W, Padilha AS, Ruddiman CA, Best AK, Macal E, Kim-Shapiro DB, Christ G, Yan Z, Cortese-Krott MM, Ricart K, Patel R, Bender TP, Sonkusare SK, Weiss MJ, Ackerman H, Columbus L, Isakson BE. Endothelial alpha globin is a nitrite reductase. Nat Commun. 2022;13(1):6405. Epub 20221027. doi: 10.1038/s41467-022-34154-3. PubMed PMID: 36302779; PMCID: PMC9613979.

62. Lechauve C, Butcher JT, Freiwan A, Biwer LA, Keith JM, Good ME, Ackerman H, Tillman HS, Kiger L, Isakson BE, Weiss MJ. Endothelial cell alpha-globin and its molecular chaperone alpha-hemoglobin-stabilizing protein regulate arteriolar contractility. The Journal of clinical investigation. 2018;128(11):5073–82. Epub 20181008. doi: 10.1172/JCI99933. PubMed PMID: 30295646; PMCID: PMC6205378.

63. Straub AC, Lohman AW, Billaud M, Johnstone SR, Dwyer ST, Lee MY, Bortz PS, Best AK, Columbus L, Gaston B, Isakson BE. Endothelial cell expression of haemoglobin alpha regulates nitric oxide signalling. Nature. 2012;491(7424):473–7. Epub 20121031. doi: 10.1038/nature11626. PubMed PMID: 23123858; PMCID: PMC3531883.

64. Opitz B, Hippenstiel S, Eitel J, Suttorp N. Extra- and intracellular innate immune recognition in endothelial cells. Thromb Haemost. 2007;98(2):319–26. PubMed PMID: 17721613.

65. Khakpour S, Wilhelmsen K, Hellman J. Vascular endothelial cell Toll-like receptor pathways in sepsis. Innate Immun. 2015;21(8):827–46. Epub 20150923. doi: 10.1177/1753425915606525. PubMed PMID: 26403174.

66. Aird WC. Endothelial cell heterogeneity. Cold Spring Harb Perspect Med. 2012;2(1):a006429. doi: 10.1101/cshperspect.a006429. PubMed PMID: 22315715; PMCID: PMC3253027.

67. Cleuren ACA, van der Ent MA, Jiang H, Hunker KL, Yee A, Siemieniak DR, Molema G, Aird WC, Ganesh SK, Ginsburg D. The in vivo endothelial cell translatome is highly heterogeneous across vascular beds. Proceedings of the National Academy of Sciences of the United States of America. 2019;116(47):23618–24. Epub 20191111. doi: 10.1073/pnas.1912409116. PubMed PMID: 31712416; PMCID: PMC6876253.

68. Jambusaria A, Hong Z, Zhang L, Srivastava S, Jana A, Toth PT, Dai Y, Malik AB, Rehman J. Endothelial heterogeneity across distinct vascular beds during homeostasis and inflammation. eLife. 2020;9. Epub 20200116. doi: 10.7554/eLife.51413. PubMed PMID: 31944177; PMCID: PMC7002042.

69. Kalucka J, de Rooij L, Goveia J, Rohlenova K, Dumas SJ, Meta E, Conchinha NV, Taverna F, Teuwen LA, Veys K, Garcia-Caballero M, Khan S, Geldhof V, Sokol L, Chen R, Treps L, Borri M, de Zeeuw P, Dubois C, Karakach TK, Falkenberg KD, Parys M, Yin X, Vinckier S, Du Y, Fenton RA, Schoonjans L, Dewerchin M, Eelen G, Thienpont B, Lin L, Bolund L, Li X, Luo Y, Carmeliet P. Single-Cell Transcriptome Atlas of Murine Endothelial Cells. Cell. 2020;180(4):764–79 e20. Epub 20200213. doi: 10.1016/j.cell.2020.01.015. PubMed PMID: 32059779.

70. Songjang W, Paiyabhroma N, Jumroon N, Jiraviriyakul A, Nernpermpisooth N, Seenak P, Kumphune S, Thaisakun S, Phaonakrop N, Roytrakul S, Pankhong P. Proteomic Profiling of Early Secreted Proteins in Response to Lipopolysaccharide-Induced Vascular Endothelial Cell EA.hy926 Injury. Biomedicines. 2023;11(11). Epub 20231115. doi: 10.3390/biomedicines11113065. PubMed PMID: 38002065; PMCID: PMC10669054.

71. Kwon OK, Lee W, Kim SJ, Lee YM, Lee JY, Kim JY, Bae JS, Lee S. In-depth proteomics approach of secretome to identify novel biomarker for sepsis in LPS-stimulated endothelial cells. Electrophoresis. 2015;36(23):2851–8. Epub 20150913. doi: 10.1002/elps.201500198. PubMed PMID: 26257168.

72. Kiger L, Keith J, Freiwan A, Fernandez AG, Tillman H, Isakson BE, Weiss MJ, Lechauve C. Redox-Regulation of alpha-Globin in Vascular Physiology. Antioxidants (Basel). 2022;11(1). Epub 20220114. doi: 10.3390/antiox11010159. PubMed PMID: 35052663; PMCID: PMC8773178.

73. Kiger L, Vasseur C, Domingues-Hamdi E, Truan G, Marden MC, Baudin-Creuza V. Dynamics of alpha-Hb chain binding to its chaperone AHSP depends on heme coordination and redox state. Biochimica et biophysica acta. 2014;1840(1):277–87. Epub 20130921. doi: 10.1016/j.bbagen.2013.09.015. PubMed PMID: 24060751.

74. Bianchi ME, Rubartelli A, Sitia R. Preferential Secretion of Oxidation-Sensitive Proteins by Unconventional Pathways: Why is This Important for Inflammation? Antioxid Redox Signal. 2024;41(10-12):693–705. Epub 20240715. doi: 10.1089/ars.2024.0554. PubMed PMID: 38916186.

75. Cruz-Garcia D, Brouwers N, Malhotra V, Curwin AJ. Reactive oxygen species triggers unconventional secretion of antioxidants and Acb1. The Journal of cell biology. 2020;219(4). doi: 10.1083/jcb.201905028. PubMed PMID: 32328640; PMCID: PMC7147093.

76. Angelini G, Gardella S, Ardy M, Ciriolo MR, Filomeni G, Di Trapani G, Clarke F, Sitia R, Rubartelli A. Antigen-presenting dendritic cells provide the reducing extracellular microenvironment required for T lymphocyte activation. Proceedings of the National Academy of Sciences of the United States of America. 2002;99(3):1491–6. Epub 20020115. doi: 10.1073/pnas.022630299. PubMed PMID: 11792859; PMCID: PMC122218.

77. Soderberg A, Sahaf B, Rosen A. Thioredoxin reductase, a redox-active selenoprotein, is secreted by normal and neoplastic cells: presence in human plasma. Cancer research. 2000;60(8):2281–9. PubMed PMID: 10786696.

78. Thomas SR, Witting PK, Drummond GR. Redox control of endothelial function and dysfunction: molecular mechanisms and therapeutic opportunities. Antioxid Redox Signal. 2008;10(10):1713–65. doi: 10.1089/ars.2008.2027. PubMed PMID: 18707220.

79. Zhao X, Alexander JS, Zhang S, Zhu Y, Sieber NJ, Aw TY, Carden DL. Redox regulation of endothelial barrier integrity. Am J Physiol Lung Cell Mol Physiol. 2001;281(4):L879–86. doi: 10.1152/ajplung.2001.281.4.L879. PubMed PMID: 11557591.

80. Kokura S, Wolf RE, Yoshikawa T, Granger DN, Aw TY. Molecular mechanisms of neutrophil-endothelial cell adhesion induced by redox imbalance. Circulation research. 1999;84(5):516–24. doi: 10.1161/01.res.84.5.516. PubMed PMID: 10082473.

